# Evolution of the nonsense mediated decay (NMD) pathway is associated with decreased cytolytic immune infiltration

**DOI:** 10.1101/535773

**Authors:** Boyang Zhao, Justin R. Pritchard

**Affiliations:** Department of Biomedical Engineering, College of Engineering, The Pennsylvania State University; Quantalarity Research Group LLC

**Keywords:** Cytolytic immune infiltration, nonsense mediated decay, immunotherapy, tumor evolution, frameshift mutations, tumor burden

## Abstract

**Background:** The somatic co-evolution of tumors and the cellular immune responses that combat them drives the diversity of immune-tumor interactions. This includes tumor mutations that generate neo-antigenic epitopes that elicit cytotoxic T-cell activity and subsequent pressure to select for genetic loss of antigen presentation. Most studies have focused on how tumor missense mutations can drive tumor immunity, but frameshift mutations have the potential to create far greater antigenic diversity. However, expression of this antigenic diversity is potentially regulated by Nonsense Mediated Decay (NMD) and NMD has been shown to be of variable efficiency in cancers.

**Methods:** Using TCGA datasets, we derived novel patient-level metrics of ‘NMD burden’ and interrogated how different mutation and most importantly NMD burdens influence cytolytic activity using machine learning models and survival outcomes.

**Results:** We find that NMD is a significant and independent predictor of immune cytolytic activity. Different indications exhibited varying dependence on NMD and mutation burden features. We also observed significant co-alteration of genes in the NMD pathway, with a global increase in NMD efficiency in patients with NMD co-alterations. Finally, NMD burden also stratified patient survival in multivariate regression models.

**Conclusions:** Our work suggests that beyond selecting for mutations that elicit NMD in tumor suppressors, tumor evolution may react to the selective pressure generated by inflammation to globally enhance NMD through coordinated amplification and/or mutation.

## Background

The co-evolutionary arms race between cancer and the immune response can drive tumor evolution. Tumors with high levels of clonal neoantigens have higher levels of T-cell infiltration^1^, and higher response rates to immunotherapies^1–3^. High levels of immune infiltration are also associated with loss of function mutations in Class 1 MHC proteins^4^, suggesting that the inflammation caused by T-cell-tumor recognition can result in selective pressure to lose T-cell tumor interactions. Many of the same variables that have been associated with immune infiltration in untreated tumors have also been associated with therapeutic response to checkpoint inhibitors^5–9^. Thus, exploring the predictors of inflammation and survival in the public TCGA datasets is an important source of hypotheses about immuno-therapeutic responses in human tumors, and gives us a window into the process of co-evolution between tumors and the immune system that shapes the immune ecology of the tumor and its microenvironment.

Only a minority of patients across multiple cancer types have been shown to be sensitive to single agent immunotherapy ^10–13^. This has prompted the rapid clinical development of anti-PD-1 antibodies alongside biomarkers in diverse patient populations and in combination with a variety of established and experimental therapeutics. Notably, combining checkpoint inhibitors with patient stratification has led to the approval of Pembrolizumab (an anti-PD-1 monoclonal antibody) in previously untreated NSCLC patients with PD-L1 positive tumors^14^. However, like targeted therapy, only a minority of NSCLC patients are PD-L1 positive. Unlike targeted therapy, patients that are PD-L1 negative have non-trivial response rates to immunotherapy.

Beyond PD-L1 positivity, other factors such as tumor mutational burden, microsatellite instability, oncogenic viruses have also been associated with therapeutic response and tumor immune infiltrates^3,4,15^. The rationale for their utility is that increased antigenic burden creates a specific T-cell response. Neoantigen burden due to non-synonymous substitutions has been clearly associated with immunotherapeutic success, cytolytic activity, and overall survival^1,2,16^. This link has also been demonstrated in a prospective clinical trial with nivolumab plus ipilimumab. In this randomized trial, the authors demonstrated that stage IV or recurrent NSCLC (not previously treated with chemotherapy and with a tumor PD-L1 expression level of less than 1%) who have more than 10 nonsynonymous mutations per megabase have a 42.6% progression free survival at 1 year^17^. However, this recent clinical trial only examined single amino acid mutations, and many prior predictions of neoantigen burden also tended to predict neoantigens using single substitution variants^16,18–20^.

Interestingly, like PD-1, many patients with high neoantigen burden fail to respond to immunotherapy, and some patients with low neoantigen burden have durable responses to immunotherapy and exhibit high levels of tumor inflammation with cytolytic cells^2,21^. As such, there is a critical need to continue to understand and predict mediators of tumor immunity in humans. Recently, multiple improvements to neoantigen predictions have been made by incorporating clonality^1^, indel mutations^22^, and intron retention mutations^23^ into studies of immune infiltration and immune response in tumors.

Specifically, frameshift mutations have been examined genomically, pre-clinically, and in patient case studies^24,25^. These revealed that mutagenic indels provide a highly immunogenic source of antigens^24^. However, deeper investigations of the different mechanisms of neoantigen generation have the potential to expand the prognostic power of clinicians to predict immunotherapy responses and indications. Moreover, better predictions of the neoantigen landscape should aid current and future efforts to develop neoantigen derived peptide vaccines. These peptide vaccines may help turn immunotherapy non-responsive tumors into immunotherapy responsive tumors.

However, the use of frameshift mutations for predictions in pan-cancer analyses raises an important question. Frameshifts were not initially considered in genomic analyses of neoantigens because they were considered to be unlikely to be expressed due to nonsense mediated decay (NMD)^1^. Frameshift mutations cause premature termination codons (PTCs), resulting in mRNAs that are the target of nonsense-mediated decay (NMD). While NMD should lead to a loss of expression of the resulting transcripts, NMD has been found to function with varying efficiency^26^. This may lead to reduced NMD and result in the expression of frameshift mutations that could be presented as neoantigens. In addition, genomic analyses of frameshift mutations and their expression suggests that NMD operates with reduced efficiency in cancer^24^. Finally, there is also evidence in preclinical models that inhibiting nonsense mediated decay can enhance tumor immunity^27^. Thus, we believe that NMD itself may also act as an independent biological filter of which indels are expressed, and thus we aimed to quantify the additional information that NMD brings to the prediction of frameshift neoantigens. We hypothesize that additional orthogonal predictive metrics of immune activation can be derived based on patient-level NMD efficiencies. Here, we examined multiple cancer indications for associations between NMD/mutational burden with cytolytic activity/survival. We find that accounting for NMD and indel mutations is significantly better than accounting for indels alone. We also find that NMD derived metrics have some independent prognostic value alongside current clinical parameters like PD-L1 expression and simple tumor mutation burden (TMB)^3,28^.

## Methods

### Datasets

The following TCGA datasets were acquired from Board Institute GDAC Firehose repository: mRNA-seq V2 RSEM level 3, mutation calls level 3, copy number level 3, and clinical data level 1, for the following indications: BLCA, BRCA, CESC, COAD/READ, GBM/LGG, HNSC, KIPAN, LUAD, LUSC, OV, PRAD, SKCM, STAD, THCA, and UCEC, downloaded over the period from October to December of 2017. MSI data was acquired from Hause et al^29^. MSI was included for analyses only for indications with at least 10 cases of MSI-H and MSS each (UCEC and STAD satisfied this criteria). Note COAD/READ also had substantial MSI-H, but the mutation calls dataset for the COAD/READ was limited (to 223 patients), thus after merging across datasets, there were too few MSI-H patients for MSI to be included for subsequent analyses.

### Data preprocessing and quality controls

The datasets were further preprocessed in preparation for analyses and modeling. For all datasets, only one tumor sample was used per patient (filter was based on sample type code). In the case of patient tumors with multiple vials, the lowest-valued vial was used. For mRNA-seq data, the transcripts per kilobase million (TPM) value was used. TPM was calculated as scaled estimate (tau value) * 1e6. For CNA data, missing CNA were treated as zero (no CNA). For calculating NMD efficiency and burden (more details in *NMD feature engineering* section), a series of data cleaning and quality controls were performed (Supp Figure S2). PTC-bearing transcripts were excluded if they overlapped with CNA. Genes were removed from the NMD efficiency and burden calculation if the WT was noisy (CV > 0.05 or < 10 samples) or low expression (median TPM < 5).

### Cytolytic activity

The cytolytic activity was calculated as the geometric mean of the expressions (in TPM+0.01) of *GZMA* and *PRF1*^4^. The cytolytic activity values were also categorized into high/low with upper/lower quartiles.

### NMD feature engineering

Metrics for nonsense-mediated decay burden was derived based on NMD efficiency values. The calculation for NMD efficiency was based on Lindeboom et al^26^. The data was preprocessed as described in the data quality controls section above. The efficiency was calculated at the gene-level, as the negative log base 2 transform of the ratio between the expression of the mutant-bearing transcript and the median mRNA expression of that transcript (calculated from samples with no mutations and copy number variations for that transcript). We accordingly derived NMD efficiencies for nonsense-, frameshift-, and nonsense/frameshift-bearing transcripts. Here it is important to clearly distinguish between NMD metrics. A measure of NMD burden is at the patient-level and is unique to this paper, while the measurement of NMD efficiency is at the gene-level and is the same as Lindeboom et al. To turn NMD efficiency measurements into estimates of NMD burden at the patient level, the NMD efficiency values were aggregated in several ways. This included calculation of the: median, mean, maximum, total number of genes with NMD, the fraction of total genes with NMD, and the median of the expression-weighted efficiency value (weighted by percentile of median WT gene expression out of all median WT gene expressions), and mean of the expression-weighted efficiency value.

### Random forest model

A random forest model was used for predicting cytolytic activity (as a binary classification of low/high) based on the engineered feature set using the *randomForest* package. The random forest is a robust machine learning model that controls for overfitting internally. The hyperparameters for our random forest models were tuned using the *caret* package across the following values: number of randomly selected variables to try at each split, mtry [Round(*α*/2), *α*, 2*α*], where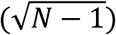 and *N* is the number of features; and number of trees to try [500, 1000]. The combination of hyperparameter values chosen for the final model was based on the model with the highest AUC. Since the overall goal of the model is to infer importance, the entire dataset with all patients was used for model building. Although random forest internally controls for overfitting, 10-fold cross-validation was also performed to check for robustness of the model. Different models were built with different subsets of feature classes (e.g. mutation, NMD burden, MSI, mutation + NMD burden, mutation + MSI, NMD burden + MSI). Pan-cancer models were also created. The individual datasets per indication were pooled together to generate the pan-cancer dataset. All metric calculations (e.g. median, etc.) were calculated at the per-indication level. Variable importance and significance were calculated using the *rfPermute* package.

### NMD alterations and pathway alterations

For NMD co-occurrence analyses, we used the cBioPortal^30^ to examine all TCGA pan-cancer datasets (pan_cancer_atlas) on Jan 8, 2019. We searched NMD genes SMG 1,5,6,7 and USP 1,2,3B. Amplifications and mutations were searched separately. For copy number alterations of different pathways, we also searched the corresponding genes in cBioPortal. Genetic alterations of NMD and their association with NMD efficiency metrics and cytolytic activity metrics were also examined, with the metrics calculated per methods described above. For copy number variations, amplifications and deep deletions were considered using copy number cutoff of 1 and −1, respectively. NMD genes with no copy number variations (copy number value of zero) and no mutations were used as controls.

### Survival analyses

Survival analyses were performed using the *survival* package in R. The survival function was estimated using Kaplan-Meier product limit method and its variance was estimated using Greenwood’s method. The hazard ratios were calculated using Cox proportional hazards model. In addition to univariate models, multivariate models were performed where each feature was controlled for age (categorical, < or ≥ 65), gender (categorical, male/female), TNM stage (categorical), TMB (categorical, ≤ or > median), and PD-L1 level (categorical, low/med/high quartiles of TPM values). TMB was determined using total mutation counts of missense, nonstop, nonsense, and frameshift divided by 38Mb, an estimate of the exome size. The p-values were corrected for multiple hypothesis testing using Benjamini-Hochberg.

### Statistical analyses

All analyses were performed in R. Correlations among the covariates were calculated with Spearman correlation. In univariate analyses of feature values, the difference between categorical cytolytic low versus high was tested using Mann-Whitney for continuous features and Chi-squared for categorical features. Statistical test for trend between NMD metrics and NMD genetic alterations was performed using Jonckheere-Terpstrata test. Statistical methods for random forest model and for survival analyses were described in their respective sections above.

### Code and data availability

Code for the analyses and outputs are available on GitHub at https://github.com/pritchardlabatpsu/NMDcyt.

## Results

### NMD metrics are orthogonal predictors of tumor cytolytic activity

Recent work by Rooney et al. used the transcript levels of two cytolytic effectors *GZMA* and *PRF1* (Supp Fig S1) to assess the immune cytolytic activity^4^. Here, using this measure for immune cytolytic activity, we quantitatively examined 17 cancer indications for the contribution of mutation variant counts to observed cytolytic activity (high versus low). We performed a pan-cancer analysis using a random forest model with the total counts of each mutation variant type as features. A final AUROC value of 0.59 suggest that using mutation counts does not fully explain the cytolytic activity, but they are statistically significant contributors (Supp Fig S2A and S2B). Almost all the mutation variants are important and contribute to the model accuracy (Supp S2C). As expected, we observed that missense, nonsense, and silent mutation variants are correlated^24^. However, frameshift mutation counts are not strongly correlated with silent mutation counts, hence suggesting frameshift are an orthogonal predictor (Supp Fig S2D).

Recent developments have suggested that frameshifts (which create very distinct neoepitopes) can improve the prediction of inflamed tumors and patient survival^24^. However, this presents a question: does patient level NMD independently associate with metrics of tumor inflammation and overall survival in a manner that is independent from indel abundance? Previous work performed an approximate correction for NMD, but, the NMD process has been shown to be complex and variable^26^, and could be measured at the patient level by many metrics. For instance, the central tendency of NMD across all transcripts should give an indication of the efficiency of the process of NMD within an individual while the maximum NMD level within an individual for a specific transcript might measure the propensity for NMD to inhibit specific neoantigens. We hypothesized that to understand the role of nonsense mediated decay more deeply, we had to investigate many measures of NMD activity simultaneously. As NMD efficiency is measured at the individual gene level, while cytolytic activity is measured at the patient-level, we began by deriving multiple patient-level measures of ‘NMD burden’, using different approaches to aggregate the NMD efficiency values (Fig 1A and Supp Fig S3). This included a burden metric of nonsense mutations (ns), frameshift mutations (fs), and combined nonsense and frameshifts (ns+fs). We first examined the correlation among the variables, and observed that related variables (i.e. NMD related metrics, cytolytic activity metrics) tended to cluster together (Fig 1B). In addition, simple metrics of mutation abundance are positively correlated with cytolytic activity while most NMD-based metrics are negatively correlated (Fig 1B, Supp Fig S4). This suggests that higher NMD efficiency lowers the expression of indels and possibly neoantigens. This is consistent with NMD suppressing neoantigens in experimental models of cancer^27^.

**Figure 1.**
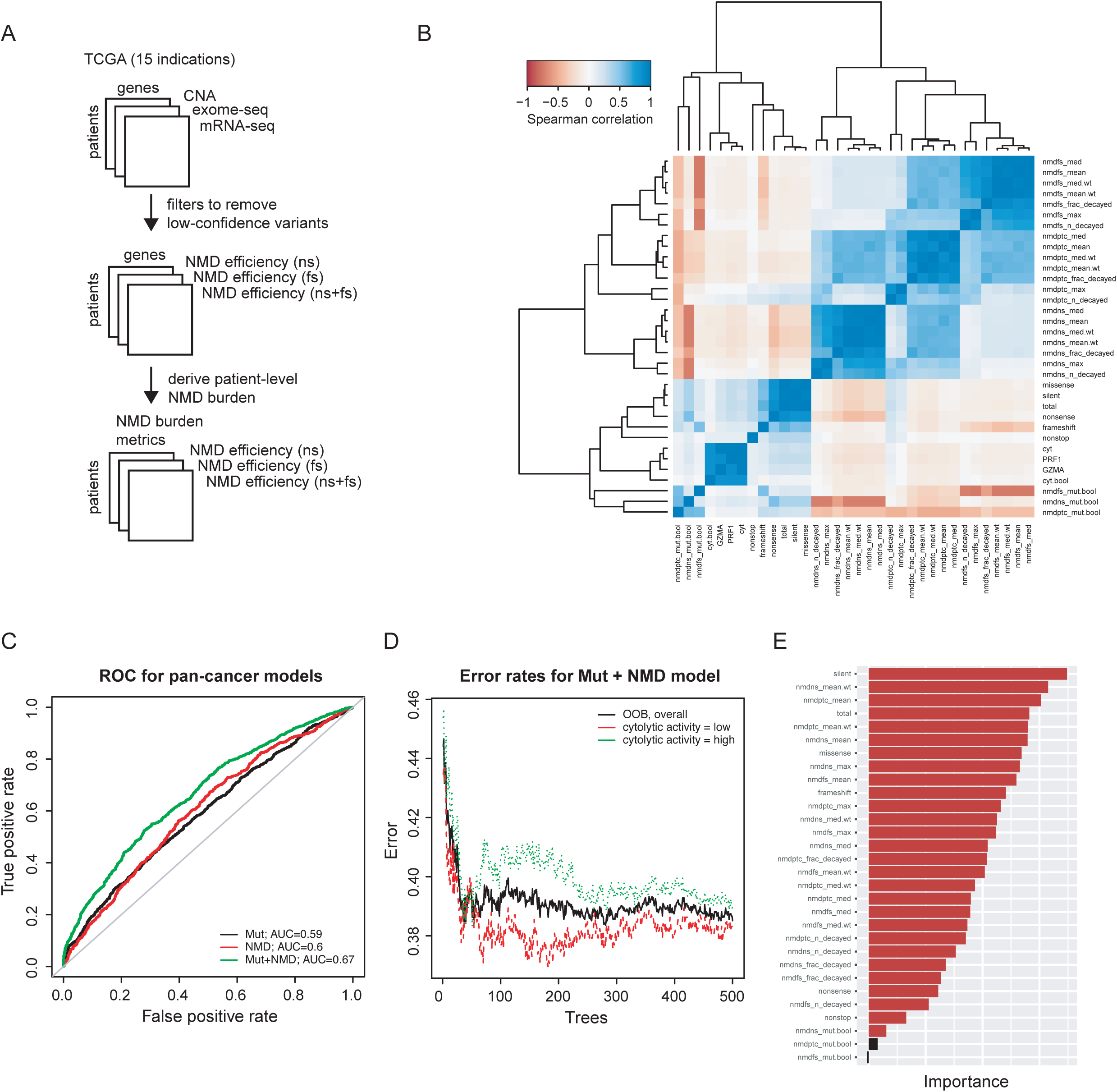
NMD burden as orthogonal predictors of cytolytic activity. (A) Schematic of data processing pipeline for deriving NMD burden, incorporating TCGA datasets for CNA, exome-seq, and mRNA-seq. (B) Pan-cancer correlation among features for mutations and NMD burden. (C) Pan-cancer ROC for random forest model with mutation variant counts only (Mut), NMD burden only (NMD), or combined (Mut+NMD). (D) Out-of-bag error of overall model (black) and for predicting cytolytic activity low (red) and high (green), for combined random forest model. (E) Variable importance of the features used in the combined model, based on mean decrease in model accuracy. nmdns: NMD metric based on nonsense transcripts; nmdfs: NMD metric based on frameshift transcripts; nmdptc: NMD metric based on nonsense and frameshift transcripts; _n_decayed: number of transcripts with NMD; _frac_decayed: fraction of transcripts with NMD; _max: maximum NMD efficiency value; _med: median NMD efficiency; _mean: mean NMD efficiency;.wt: NMD efficiency metric weighted by mRNA expression;

Using our best NMD features which tended to be measures of central tendency, we built pan-cancer models using mutation counts only, NMD-burden only, or a combination of the two feature groups. Surprisingly, NMD alone was as good of a predictor of pan-cancer cytolytic activity as mutations (AUROCs of ∼0.6). Importantly, the combined model improved upon single data type predictors with an AUROC of 0.67 (Fig 1C and 1D). Thus, mutation counts and NMD-burden offer equally important orthogonal information that combines to improve our understanding of cytolytic activity in tumors (Fig 1D and 1E).

### The NMD pathway is significantly and co-ordinately altered in cancer

A potential explanation for an association between the central tendency of NMD across all genes within a patient and cytolytic activity is that T-cell infiltration might exert selective pressure upon tumors to mutate or amplify genes in the NMD pathway. Towards this hypothesis we utilized the TCGA pan-cancer data sets. Focusing on the major genes in the NMD pathway (SMG1,5,6,7 and UPF1,2,3B)^31^, we first examined whether these genes were amplified in cancer. Examining all patients in the Pan-Cancer dataset, we observed amplification of NMD pathway genes (Fig 2A) that resembled a gain of function pathway such as MAPK family members more than they did tumor suppressors such as P53 and PTEN (i.e. more amplifications than deletions were observed, Supp Fig S5). Interestingly, across these 7 genes, all permutations of the pairwise interactions between the 7 genes co-occurred more often than one would expect by chance (corrected p-values<0.05, Fig 2B). Only one of these pairwise mutual amplifications, SMG5-SMG7, contains 2 genes that reside in a similar genomic location on chromosome 1. Beyond amplification, while none of the individual genes in the NMD pathway are predicted to be drivers of cancer, we thought it was possible that tumor evolution might select for co-occurrence of multiple individual NMD pathway mutations. Surprisingly, all 21 pairwise combinations of the 7 core NMD pathway genes exhibited a tendency towards mutational co-occurrence at a multiple hypothesis corrected p-value <0.001 (Fig 2B). Importantly, this co-alteration appears to also have a functional consequence. When we examined patients with no NMD alterations versus patients with co-alterations (i.e. >1 alteration) we observed a significant trend towards increasing NMD efficiency as NMD genes became coordinately altered (Fig 2C and Supp Fig S6 and S7). Moreover, patients with NMD alterations had lower cytolytic activity (Supp Fig S8).

**Figure 2.**
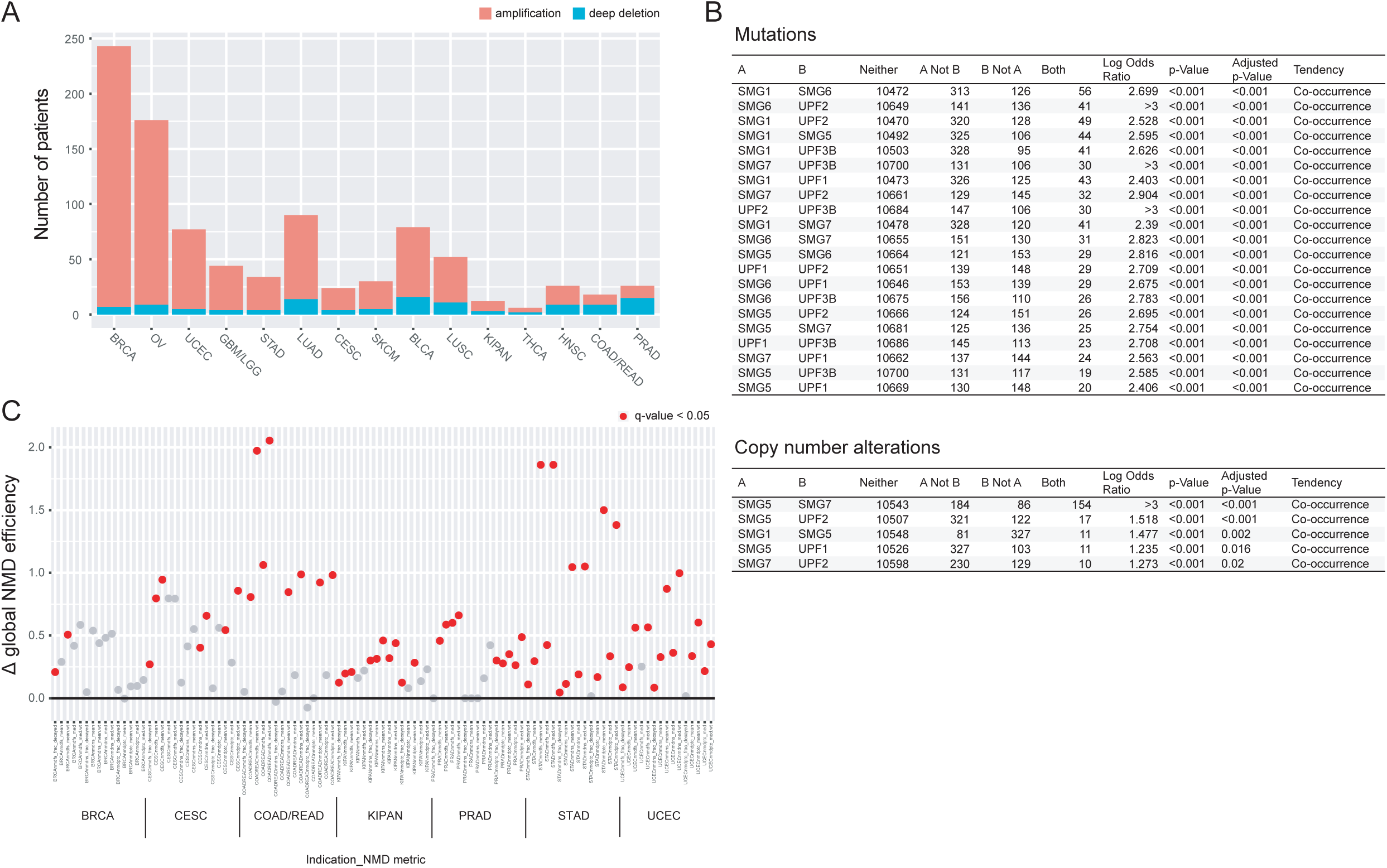
NMD alterations co-occur and associated with improved NMD efficiency. (A) Amplifications/deletions of genes in the NMD pathway (SMG1,5,6,7 and UPF1,2,3B) across different indications. (B) Co-occurrence of copy number and mutations of NMD genes, using the TCGA pan-cancer atlas datasets on cBioPortal. (C) NMD efficiency of patients with co-altered NMD genes versus those without any alterations. Y-values are shown as the difference in median of log10 transformed NMD metric values (co-altered versus no alterations). Dots shown in red are statistically significant with adjusted p-value < 0.05; Mann-Whitney test with Benjamini-Hochberg multiple hypothesis correction.

### Effects of NMD burden in individual cancer types

We next examined the contribution of our NMD metrics to predicting cytolytic activity in each individual indication in the TCGA. We observed varying AUROC patterns of mutations, counts, NMD burden, and combined models across the different indications (Fig 3 and Table S1). Notably, for example, the contribution of NMD burden in GBM/LGG and LUSC toward predicting cytolytic activity were minimal while mutational counts added predictive value. On the other hand, in BLCA and LUAD, NMD burden contributed more than mutational counts toward predicting cytolytic activity. In indications COAD/READ, SKCM, HNSC, KIPAN, STAD, and UCEC, both NMD and mutational burden were important. In 6 out of 15 indications, the combined model resulted in AUROC values better than the individual models. We next examined the individual prediction metrics within each feature category across the indications to infer which processes were important in which tumor type. The metric values varied in significance across all the TCGA indications (Supp Fig S9) with varying levels of contribution/significance toward the final model (Supp Fig S10). The features that contributed toward multiple indications included counts of missense, silent, and total mutations, and several of the NMD burden metrics (mostly as measured by mean or median) (Supp Fig S10). Most importantly, we observed that the metrics contributing toward the model are concordant with the final AUROC values of the model (e.g. indications where only NMD metrics are important based on AUROC values (Fig 3) also had only NMD metrics as important and statistically significant (Supp Fig S10)).

**Figure 3.**
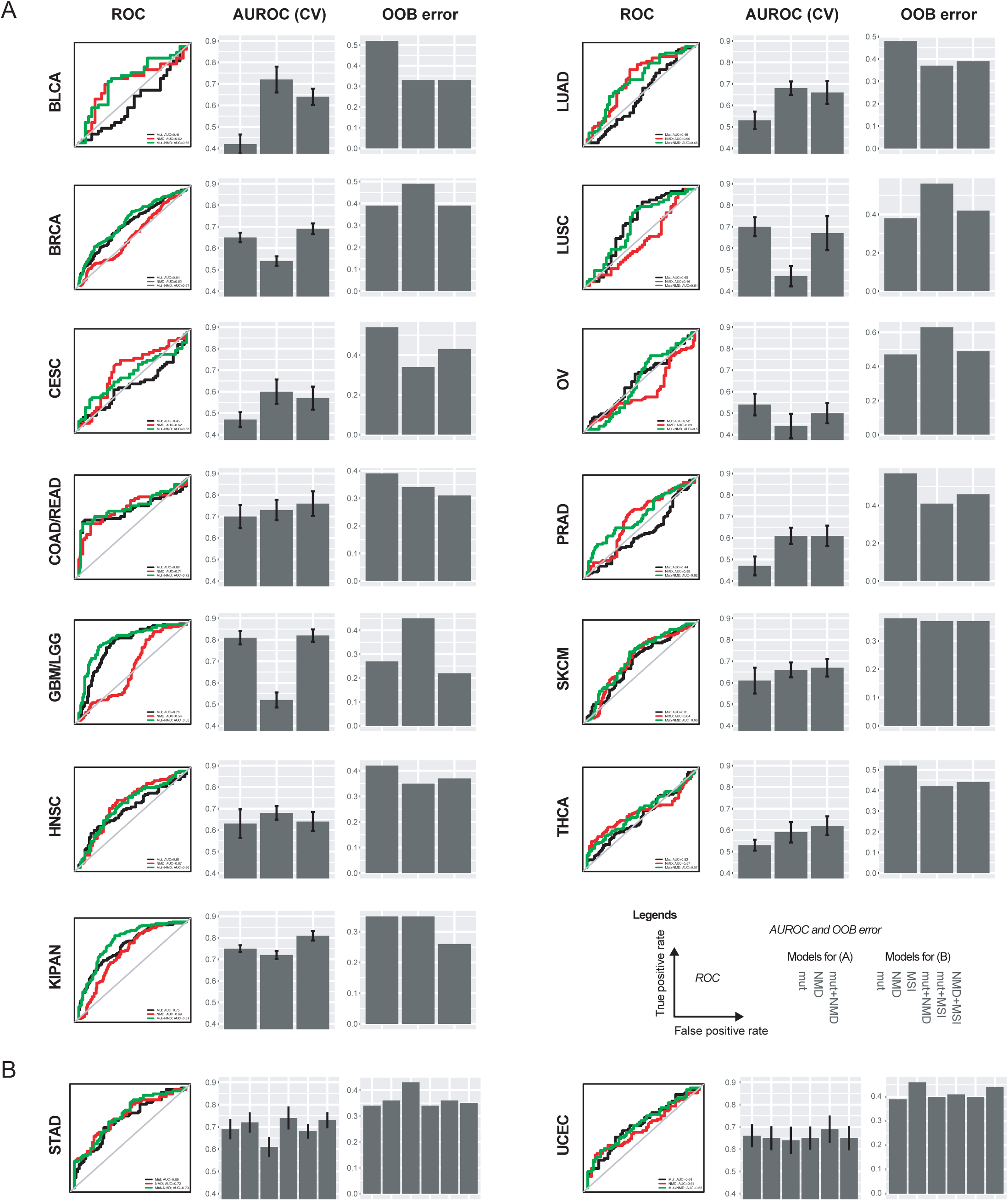
NMD burden improves predictivity of cytolytic activity. Individual ROC, AUROC, and out-of-bag (OOB) error of random forest models, for (A) indications without MSI incorporated into model, and (B) indications with MSI incorporated into model. AUROC data are shown as AUROC ± SE of AUROC from cross-validation.

### Effects of NMD burden on survival outcomes

In addition to cytolytic activity, we also examined the effects of NMD burden on survival. Here, we used the overall survival from TCGA clinical data and performed univariate Cox regression for each feature across all the indications (Supp Table S2). In melanoma for example, as expected, higher cytolytic activity is associated with a better survival outcome (hazard ratio of cyt-high vs cyt-low, 0.5; 95% CI, 0.3-0.7); p-value=0.002) (Fig 4A). A NMD burden metric was also shown to contribute to the overall survival differences. Here a high NMD burden leads to lower cytolytic activity and a worse overall survival (Fig 4B). We have also examined other known contributors, including tumor mutational burden (TMB) and PD-L1 levels, both of which were also found to stratify the survival outcomes (Supp Fig S11). Upon controlling for the covariates TMB, PD-L1, age, gender, and TNM stage, we again examined the NMD burden metric and found it to be significant. In the multivariate model, the statistical significant variables included nmdptc_med, age, gender, TMB and PD-L1 (Fig 4C).

**Figure 4.**
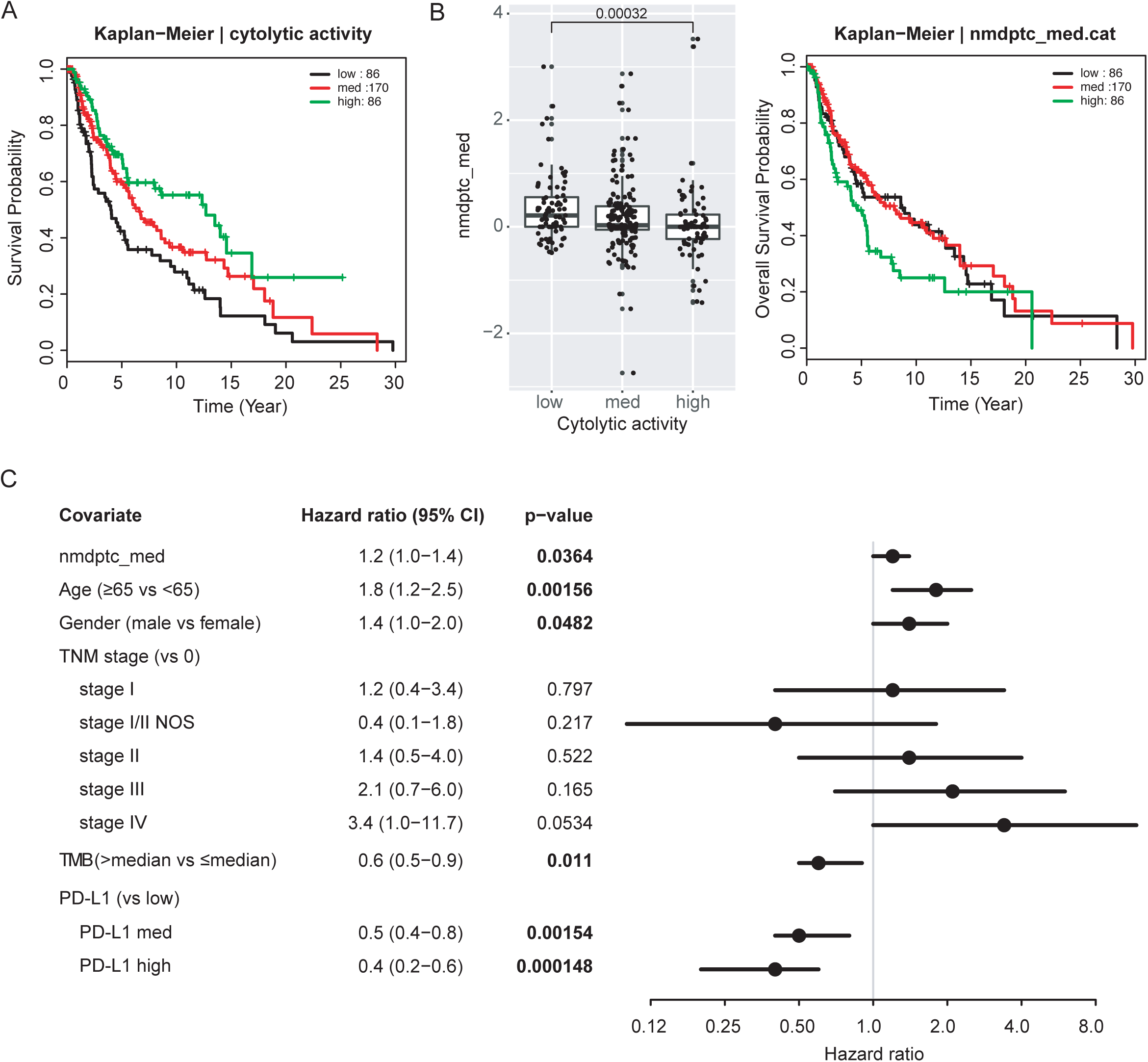
Cytolytic activity and NMD burden stratifies overall survival outcomes. (A) Kaplan-Meier overall survival for SKCM, stratified by cytolytic activity (categorized into low, med, and high based on quartiles). Legend shows the number of patients in each cytolytic activity level. (B) Association of feature nmdptc_med (NMD burden based on nonsense and frameshift, calculated based on median) on cytolytic activity. Statistical significance was determined using Mann-Whitney test. (C) Kaplan-Meier overall survival for SKCM, stratified by nmdptc_med (categorized into low, med, and high based on quartiles). (D) Forest plot of hazard ratios of each variable, in a multivariate Cox regression model for NMD burden (nmdptc_med), controlling for age, gender, TNM stage, tumor mutational burden (TMB), and PD-L1 levels.

## Discussion

A tumor’s mutational burden/neoantigen repertoire has been associated with inflammation, overall survival and therapeutic response to immunotherapies. Previous studies in the TCGA and in patients treated with checkpoint inhibitors have identified similar variables that predict immune infiltration and checkpoint response^4,8,32–34^. Because frameshift neoantigen availability is hypothetically regulated by the efficiency of the NMD process, and functional evidence has suggested that inhibition of NMD can induce a tumor immune response and tumor regression in pre-clinical models by exposing neoantigens^27^, we derived orthogonal patient-level metrics based on gene level NMD efficiencies^26^ that improved our ability to predict tumor cytolytic activity both within and across TCGA cancer types. Tumors with co-amplifications and mutations in the NMD pathway had increased NMD efficiencies and were less likely to have immune infiltration. Beyond cytolytic activity we found that NMD metrics stratified some cancer types by distinct overall survival outcomes. These stratifications were significant, even when 2 established clinical covariates PDL1 status and TMB were included.

In tumor suppressors, mutations arise spontaneously, and if they occur in a region that elicits NMD, the mutation can be selected for because NMD will eliminate the mRNA of the tumor suppressor. Clearly tumor evolution does not need to alter NMD to create loss of function in tumor suppressor proteins, it is simply selecting for mutations that happen to elicit NMD. However, our study suggests that at the level of a patient (and not a gene), mutations in NMD and a global increase in NMD efficiency can be selected for. The origin of the alterations in the NMD process could be due to enhanced suppression of tumor suppressor genes, or enhanced suppression of tumor neoantigens. Regardless of the causative selective pressure, the consequence is a tendency for genomic amplifications and mutations to co-occur within the key proteins in the NMD machinery (SMG 1,5,6,7, and UPF 1,2,3B). Consistent with this co-alteration, we observed that these co-occurring mutations/amplifications increased NMD efficiency in patients that had low cytolytic activity. This suggests that pan-cancer tumor evolution might select for co-alteration in the NMD pathway. The association of amplifications and not deletions as well as the observed functional increase in NMD efficiency suggests that cytolytic activity is inhibited by gain of function alterations in the NMD pathway.

The identification of correlates of tumor immune activity is a broad field with numerous potential candidates. One difficulty in interpreting these studies is that many studies find significant associations without quantifying the proportion of the data explained (and unexplained) by those variables. Thus, deciding which covariates to add, and how to quantify progress towards full prediction of cytolytic activity is difficult without the careful presentation of how much we understand versus how much we do not. Therefore, we present ROC curves and out-of-bag error estimates for all models. This quantitation allows us to clearly see that while we have identified significant predictors, our AUCs are mostly between 0.6 and 0.7, except for the pan-kidney dataset that has an AUC >0.8.

A potential weakness of this study is the use of mutation burden rather than MHC class I binding predictions. However, predicting neoantigen-MHC class I binding remains challenging and prune to high false positives^35^. We chose here to instead focus on mutation burden to minimize confounders from imperfect predictions. In addition, previous studies showed that mutation burden is highly correlated with neoantigen burden^4,24^, and as such likely to harbor similar information.

Beyond predictions of cytolytic activity, we quantified tumor mutation burden^36^ and PD-L1 positivity across all sample and indications in the TCGA. In melanoma the effect of NMD significantly added to a multivariate Cox regression model. Though the effect was more modest after correcting for co-variates, we suggest that it is easy to examine NMD in future clinical datasets, and as such we recommend that groups performing biomarker studies in treated and untreated patients should add metrics of NMD to attempt to understand and stratify responses.

In addition to better understanding immunity and, potentially, the response to checkpoint therapy, the inhibition of Nonsense Mediated Decay (NMD) has been proposed as a therapeutic strategy. Suppression of NMD can create a therapeutic effect through cell intrinsic (via restoring the activity of a tumor suppressor) or extrinsic mechanisms (via tumor immunity). Should a clinical candidate to inhibit NMD arise, the road to biomarker driven application of the cell intrinsic therapeutic effects is clear, but picking indications for the immune dependent activity requires studies like this one. Thus, we suggest that indications whose cytolytic activity is particularly well explained by NMD, or patients with alterations in the NMD pathway that increase NMD efficiencies might be interesting indications to look for cell non-autonomous immune driven efficacy of future NMD inhibitors.

## Conclusions

We have described here that tumor evolution may select via coordinated genetic alterations to globally enhance NMD efficiency. This in turn can influence tumor-immune interactions as measured by cytolytic activity and ultimately patient outcome.

## Abbreviations

NMD: nonsense-mediated decay
TMB: tumor mutational burden
MHC: major histocompatibility complex
PTC: premature termination codons

## Acknowledgements

We thank Pritchard lab members for helpful comments and criticism on the data and presentation.

## Availability of data and materials

Source code, outputs, and input datasets are all available through GitHub at https://github.com/pritchardlabatpsu/NMDcyt.

## Authors’ contributions

BZ and JP designed experiments, performed computational analyses, and analyzed data. BZ and JP wrote the manuscript.

## Ethics approval and consent to participate

Not applicable

## Consent for publication

Not applicable

