## Supplemental Figures for "Evolution of the nonsense mediated decay (NMD) pathway is associated with decreased cytolytic immune infiltration"

**Supplemental figure legends**

**Figure S1** Mutation variant counts are associated with cytolytic activity, albeit small. (A) Pan-cancer ROC curve for predicting cytolytic activity, using a random forest model with only counts of each mutation variant type (B) Out-of-bag error of overall model (black) and for predicting cytolytic activity low (red) and high (green). (C) Variable importance of the features used in the model, based on mean decrease in model accuracy. (D) Association in mutation counts among different mutation variant types. Missense, silent, and nonsense are correlated while frameshift is not.

**Figure S2** Correlation of gene expressions for *GZMA* and *PRF1*, component genes used for cytolytic activity.

**Figure S3** Pipeline schematic for data preprocessing and metric calculations for NMD burden. The mRNA-seq, CNA, and exome-seq datasets were incorporated. Noisy genes were filtered out, followed by derivation of gene-level NMD efficiency values. expr, expression.

**Figure S4** Univariate regression of each feature to cytolytic activity, aggregated by indication. Features were grouped into mutations (A), NMD frameshift-bearing (fs) (B), NMD nonsense-bearing (ns) (C), and NMD nonsense/frameshift-bearing (ptc) (D).

**Figure S5** Copy number alterations in different pathways across multiple indications, using the TCGA pan-cancer atlas datasets on cBioPortal. Amplifications are shown in red and deletions in blue.

**Figure S6** NMD efficiency of patients with co-amplified NMD genes versus those without any alterations. Y-values are shown as the difference in median of log10 transformed NMD metric values (co-altered versus no alterations). Dots shown in red are statistically significant with adjusted p-value < 0.05; Mann-Whitney test with Benjamini-Hochberg multiple hypothesis correction.

**Figure S7** Association between NMD co-alterations and global NMD efficiency (measured by different patient-level NMD metrics). Statistical significance was assessed with Jonckheere-Terpstrata test for trends and multiple hypothesis corrected using Benjamini-Hochberg method. Results with adjusted p-value < 0.05 are shown. The p-values shown in the plots are based on Mann-Whitney test of pair-wise comparisons.

**Figure S8** Association between NMD co-alterations and cytolytic activity. Statistical significance was assessed with Mann-Whitney test and multiple hypothesis corrected using Benjamini-Hochberg method. Results with adjusted p-value < 0.05 are shown. Nominal p-values shown in plots.

**Figure S9** Distribution of values for each feature per indication. Features were grouped into mutations (A), NMD frameshift-bearing (fs) (B), NMD nonsense-bearing (ns) (C), and NMD nonsense/frameshift-bearing (ptc) (D).

**Figure S10** Random forest model (with mutation variant counts (Mut) and NMD burden (NMD) combined) feature summaries for all indications. (A) Mean decrease in model accuracy when a given feature is removed from the model. (B) Standardized mean decease in accuracy values from (A). (C) Statistically significant features from the model, with p-value < 0.05 marked in black.

**Figure S11** Univariate overall survival analysis of SKCM for TMB (A) and PDL1 (B).

Figure S1

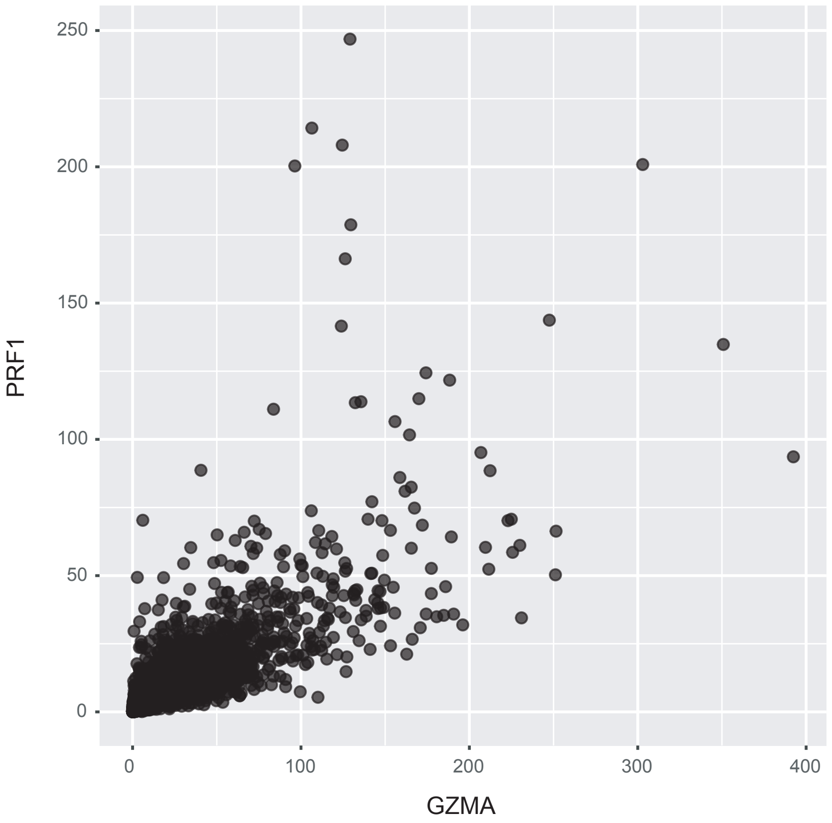

Figure S2

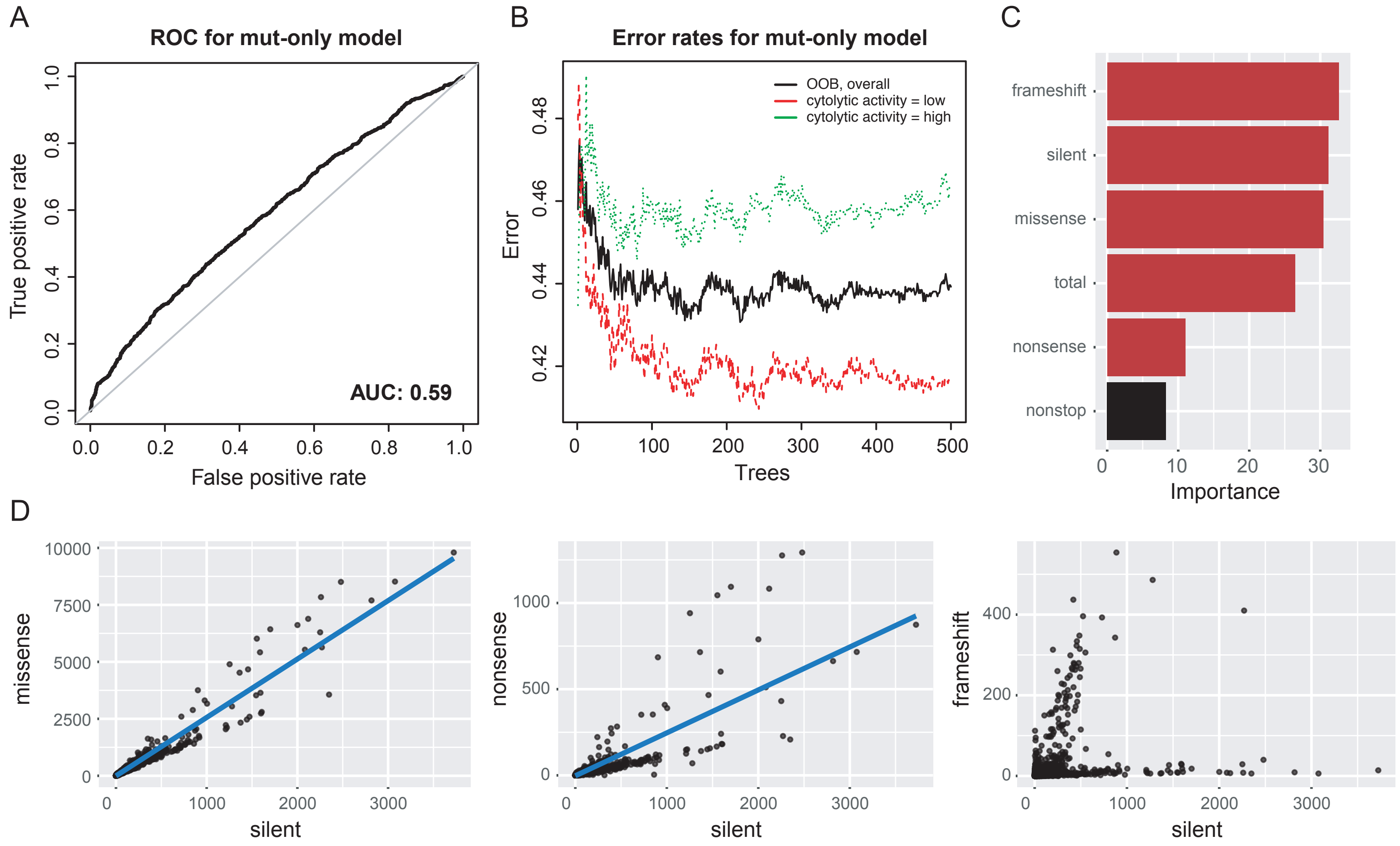

A
ROC for mut-only model
True positive rate
False positive rate
AUC: 0.59
B
Error rates for mut-only model
OOB, overall
cytolytic activity = low
cytolytic activity = high
Error
Trees
C
frameshift
silent
missense
total
nonsense
nonstop
Importance
D
missense
nonsense
frameshift
silent

Figure S3

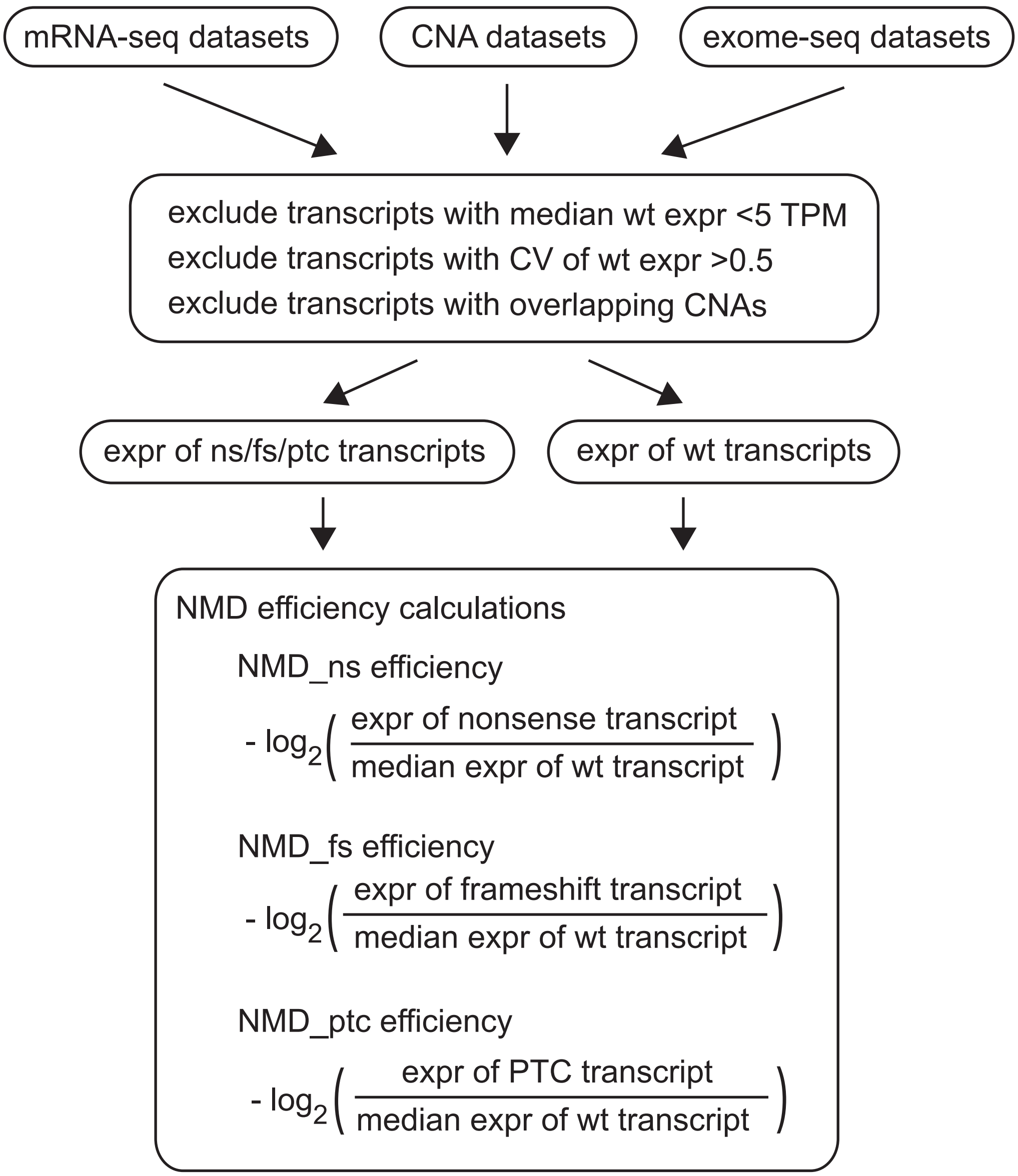
mRNA-seq datasets
CNA datasets
exome-seq datasets
exclude transcripts with median wt expr <5 TPM
exclude transcripts with CV of wt expr >0.5
exclude transcripts with overlapping CNAs
expr of ns/fs/ptc transcripts
expr of wt transcripts
NMD efficiency calculations
NMD_ns efficiency
- log2( expr of nonsense transcript / median expr of wt transcript )
NMD_fs efficiency
- log2( expr of frameshift transcript / median expr of wt transcript )
NMD_ptc efficiency
- log2( expr of PTC transcript / median expr of wt transcript )

Figure S4

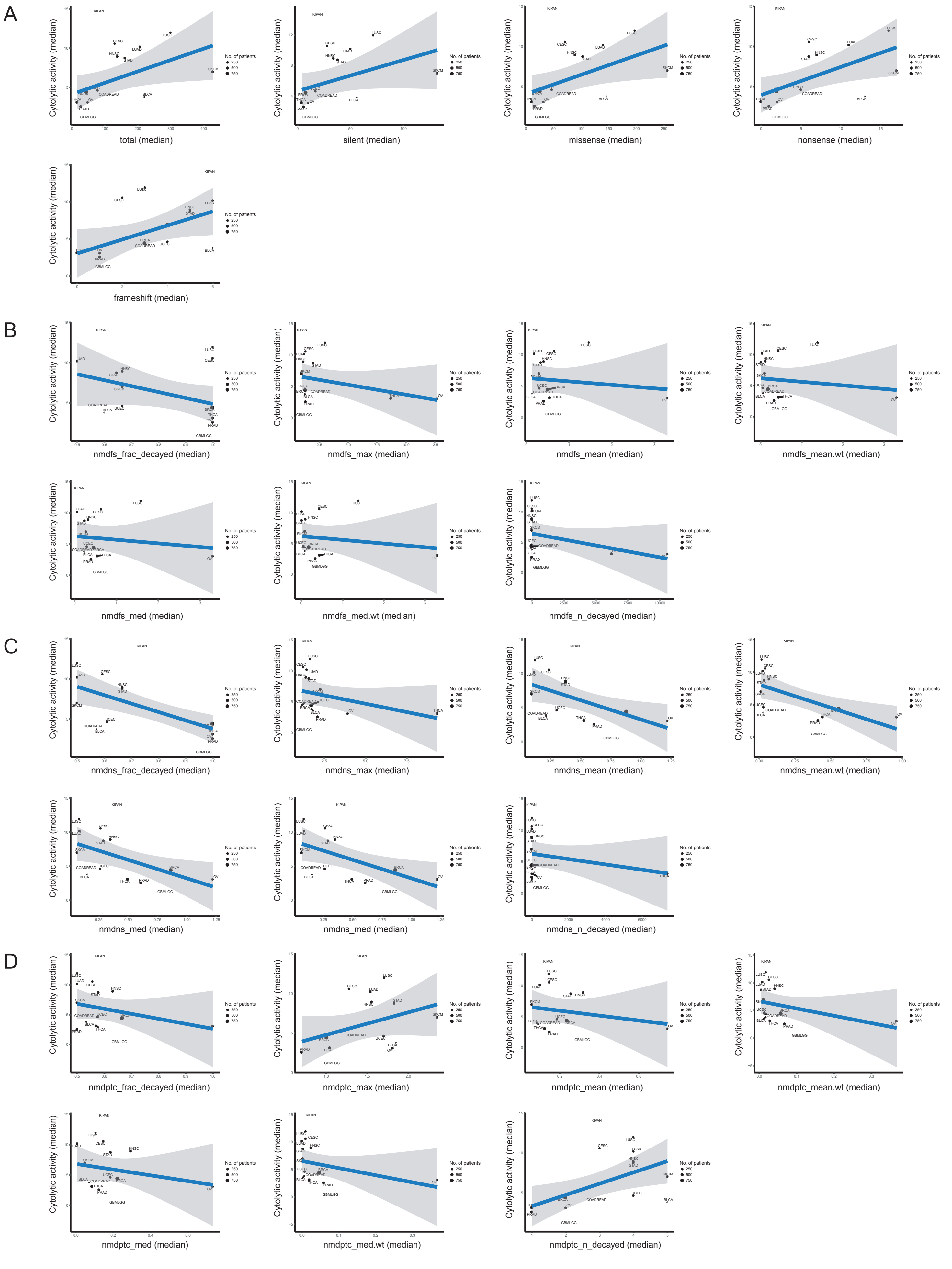
A
B
C
D
Cytolytic activity (median)
No. of patients
250
500
750
total (median)
silent (median)
missense (median)
nonsense (median)
frameshift (median)
nmdfs_frac_decayed (median)
nmdfs_max (median)
nmdfs_mean (median)
nmdfs_mean.wt (median)
nmdfs_med (median)
nmdfs_med.wt (median)
nmdfs_n_decayed (median)
nmdns_frac_decayed (median)
nmdns_max (median)
nmdns_mean (median)
nmdns_mean.wt (median)
nmdns_med (median)
nmdns_med (median)
nmdns_n_decayed (median)
nmdptc_frac_decayed (median)
nmdptc_max (median)
nmdptc_mean (median)
nmdptc_mean.wt (median)
nmdptc_med (median)
nmdptc_med.wt (median)
nmdptc_n_decayed (median)

Figure S5

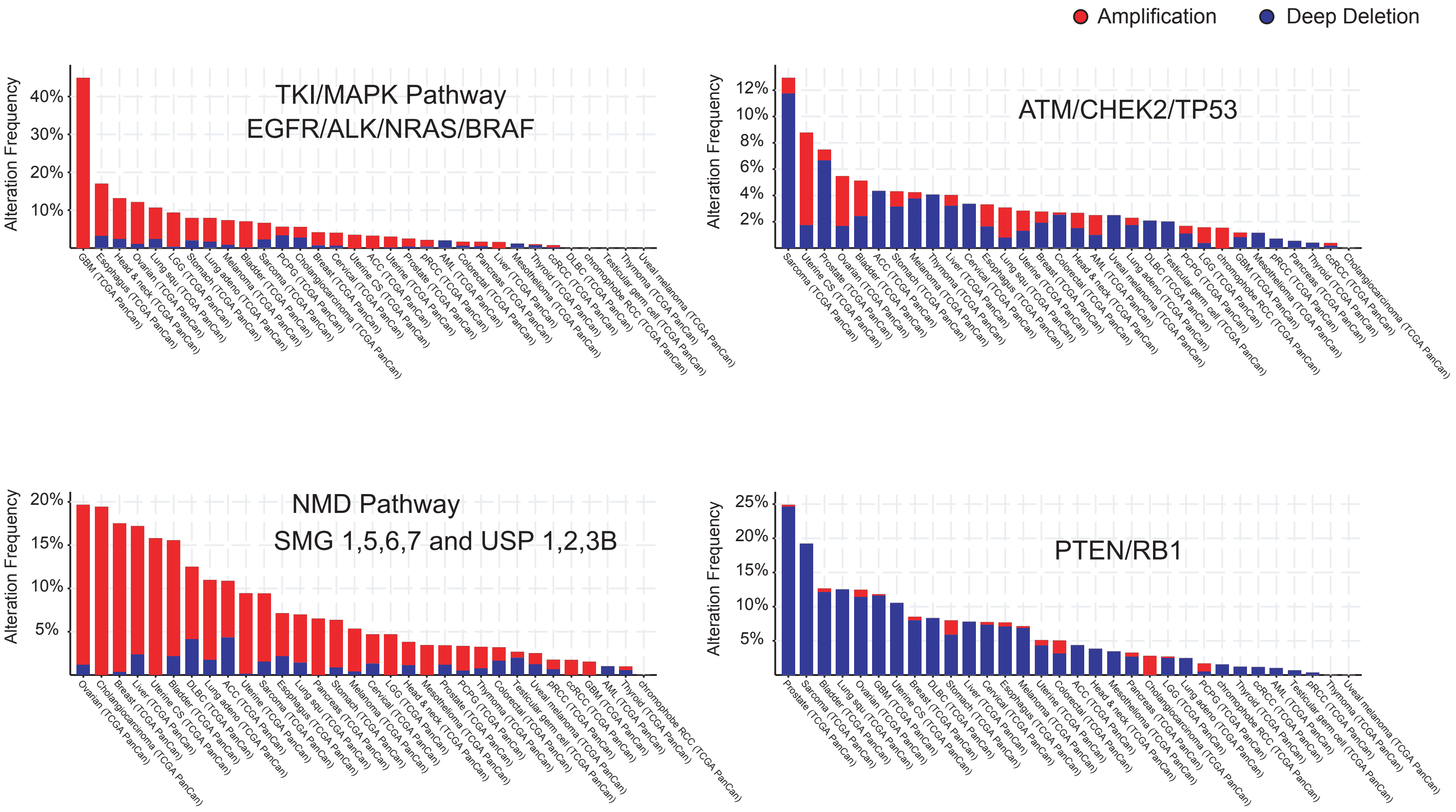
Amplification
Deep Deletion
TKI/MAPK Pathway
EGFR/ALK/NRAS/BRAF
Alteration Frequency
40%
30%
20%
10%
ATM/CHEK2/TP53
Alteration Frequency
12%
10%
8%
6%
4%
2%
NMD Pathway
SMG 1,5,6,7 and USP 1,2,3B
Alteration Frequency
20%
15%
10%
5%
PTEN/RB1
Alteration Frequency
25%
20%
15%
10%
5%

Figure S6

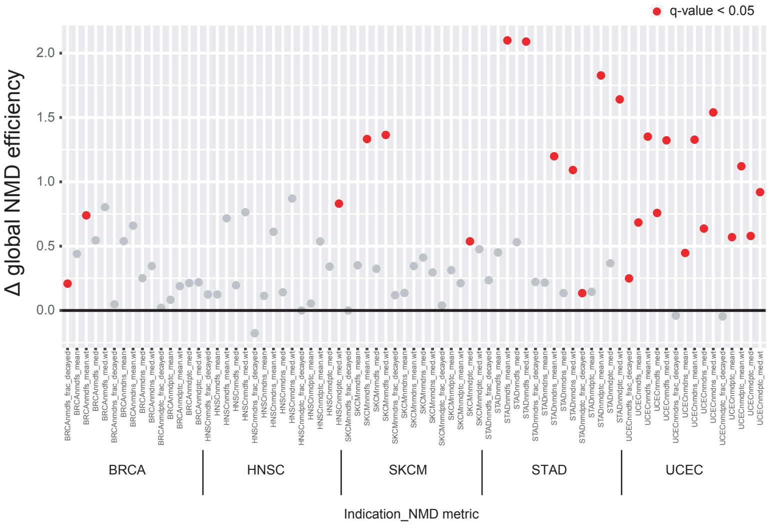

q-value < 0.05
Δ global NMD efficiency
2.0
1.5
1.0
0.5
0.0
BRCA
HNSC
SKCM
STAD
UCEC
Indication_NMD metric

Figure S7

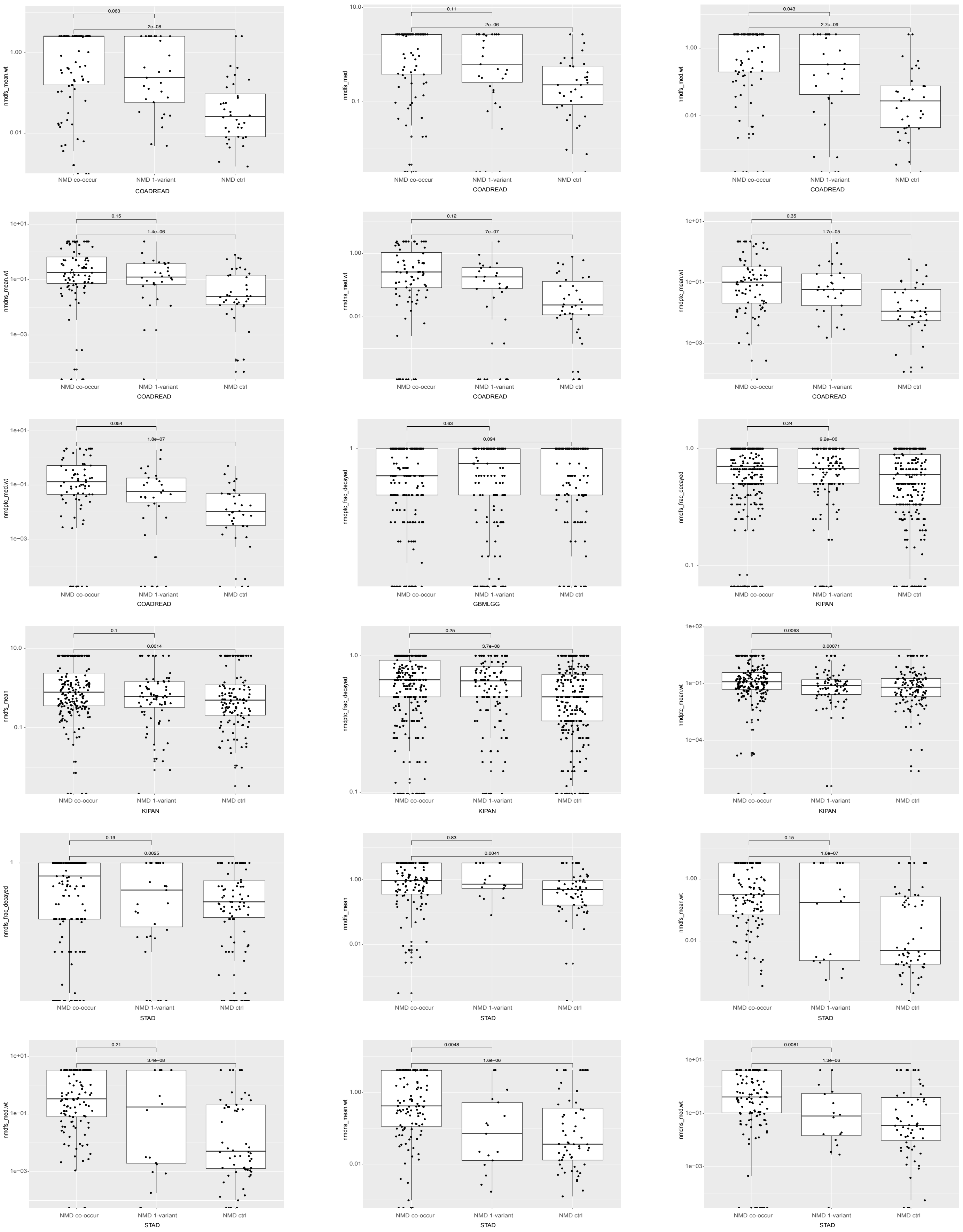
0.063
2e−08
nmdfs_mean.wt
NMD co-occur
NMD 1-variant
NMD ctrl
COADREAD
0.11
2e−06
nmdfs_med
COADREAD
0.043
2.7e−09
nmdfs_med.wt
COADREAD
0.15
1.4e−06
nmdns_mean.wt
COADREAD
0.12
7e−07
nmdns_med.wt
COADREAD
0.35
1.7e−05
nmdptc_mean.wt
COADREAD
0.054
1.8e−07
nmdptc_med.wt
COADREAD
0.63
0.094
nmdptc_frac_decayed
GBMLGG
0.24
9.2e−06
nmdfs_frac_decayed
KIPAN
0.1
0.0014
nmdns_mean
KIPAN
0.25
3.7e−08
nmdptc_frac_decayed
KIPAN
0.0063
0.00071
nmdptc_mean.wt
KIPAN
0.19
0.0025
nmdns_frac_decayed
STAD
0.83
0.0041
nmdns_mean
STAD
0.15
1.6e−07
nmdfs_mean.wt
STAD
0.21
3.4e−08
nmdns_med.wt
STAD
0.0048
1.6e−06
nmdns_mean.wt
STAD
0.0081
1.3e−06
nmdns_med.wt
STAD

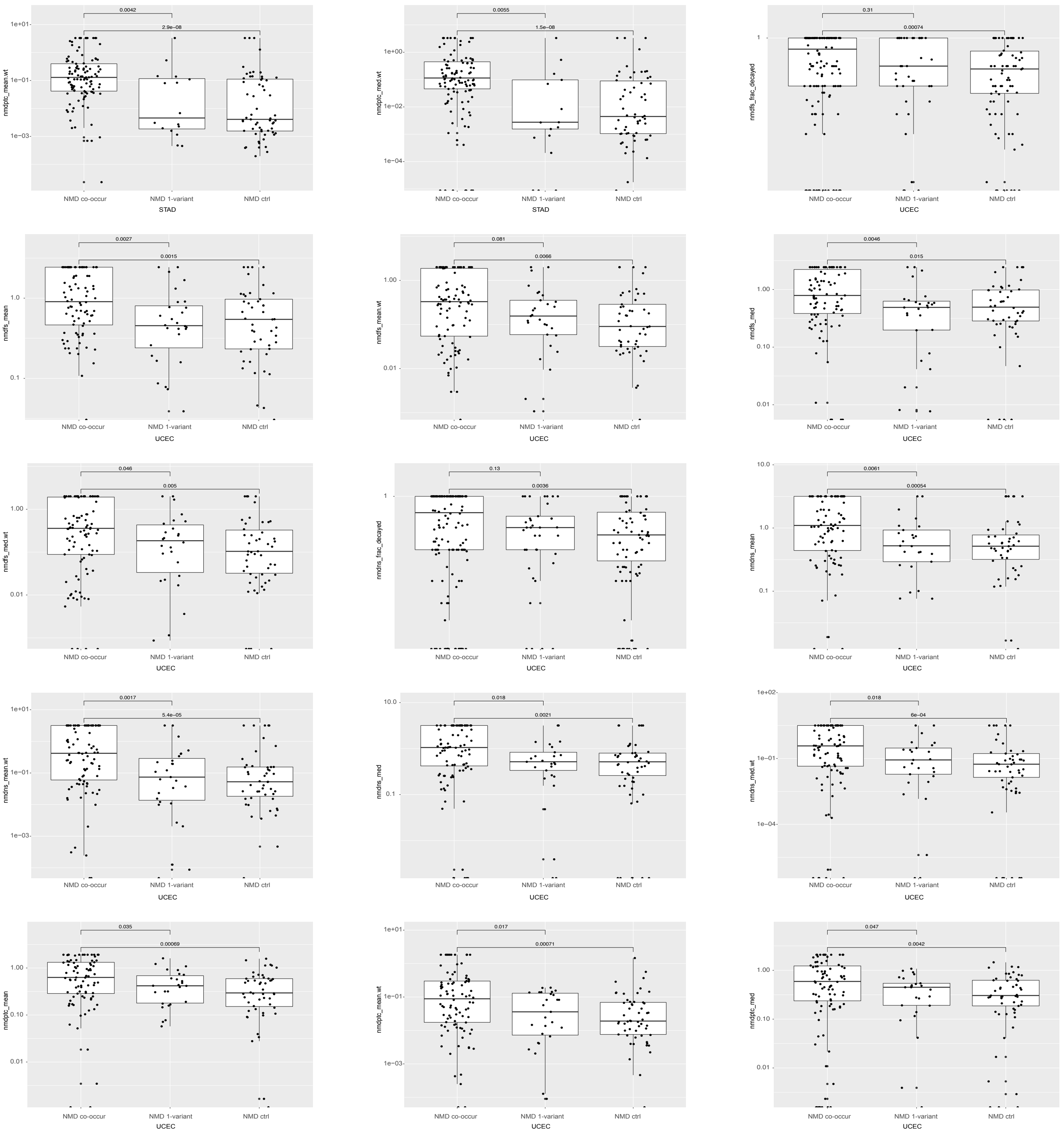

nmdptc_mean.wt
0.0042
2.9e-08
NMD co-occur
NMD 1-variant
NMD ctrl
STAD
nmdptc_med.wt
0.0055
1.5e-08
STAD
nmdfs_frac_decayed
0.31
0.00074
UCEC
nmdfs_mean
0.0027
0.0015
UCEC
nmdfs_mean.wt
0.081
0.0066
UCEC
nmdfs_med
0.0046
0.015
UCEC
nmdfs_med.wt
0.046
0.005
UCEC
nmdns_frac_decayed
0.13
0.0036
UCEC
nmdns_mean
0.0061
0.00054
UCEC
nmdns_mean.wt
0.0017
5.4e-05
UCEC
nmdns_med
0.018
0.0021
UCEC
nmdns_med.wt
0.018
6e-04
UCEC
nmdptc_mean
0.035
0.00069
UCEC
nmdptc_mean.wt
0.017
0.00071
UCEC
nmdptc_med
0.047
0.0042
UCEC

Figure S8

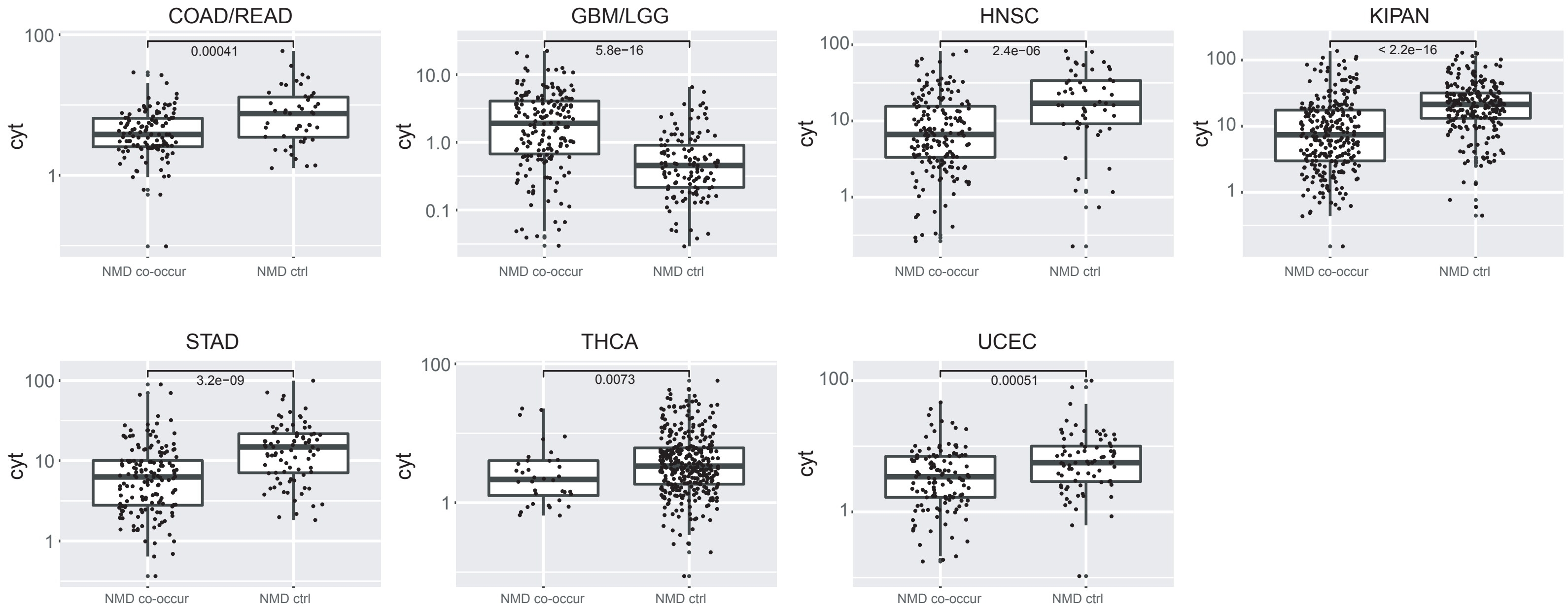
COAD/READ
0.00041
GBM/LGG
5.8e−16
HNSC
2.4e−06
KIPAN
< 2.2e−16
STAD
3.2e−09
THCA
0.0073
UCEC
0.00051
cyt
NMD co-occur
NMD ctrl
100
10
1
10.0
1.0
0.1

Figure S9

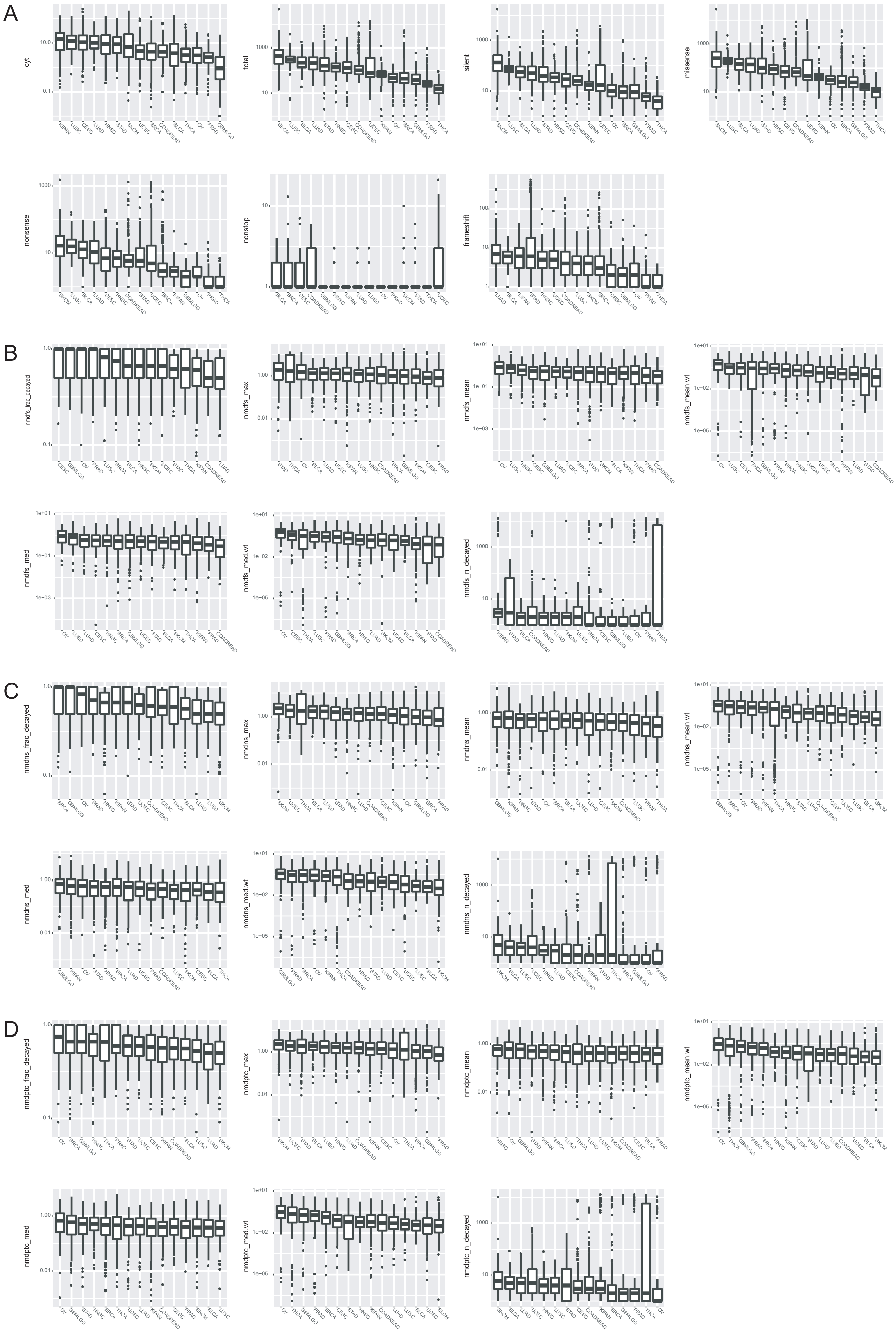
A
cyt
total
silent
missense
nonsense
nonstop
frameshift
B
nmdfs_frac_decayed
nmdfs_max
nmdfs_mean
nmdfs_mean.wt
nmdfs_med
nmdfs_med.wt
nmdfs_n_decayed
C
nmdns_frac_decayed
nmdns_max
nmdns_mean
nmdns_mean.wt
nmdns_med
nmdns_med.wt
nmdns_n_decayed
D
nmdptc_frac_decayed
nmdptc_max
nmdptc_mean
nmdptc_mean.wt
nmdptc_med
nmdptc_med.wt
nmdptc_n_decayed

Figure S10

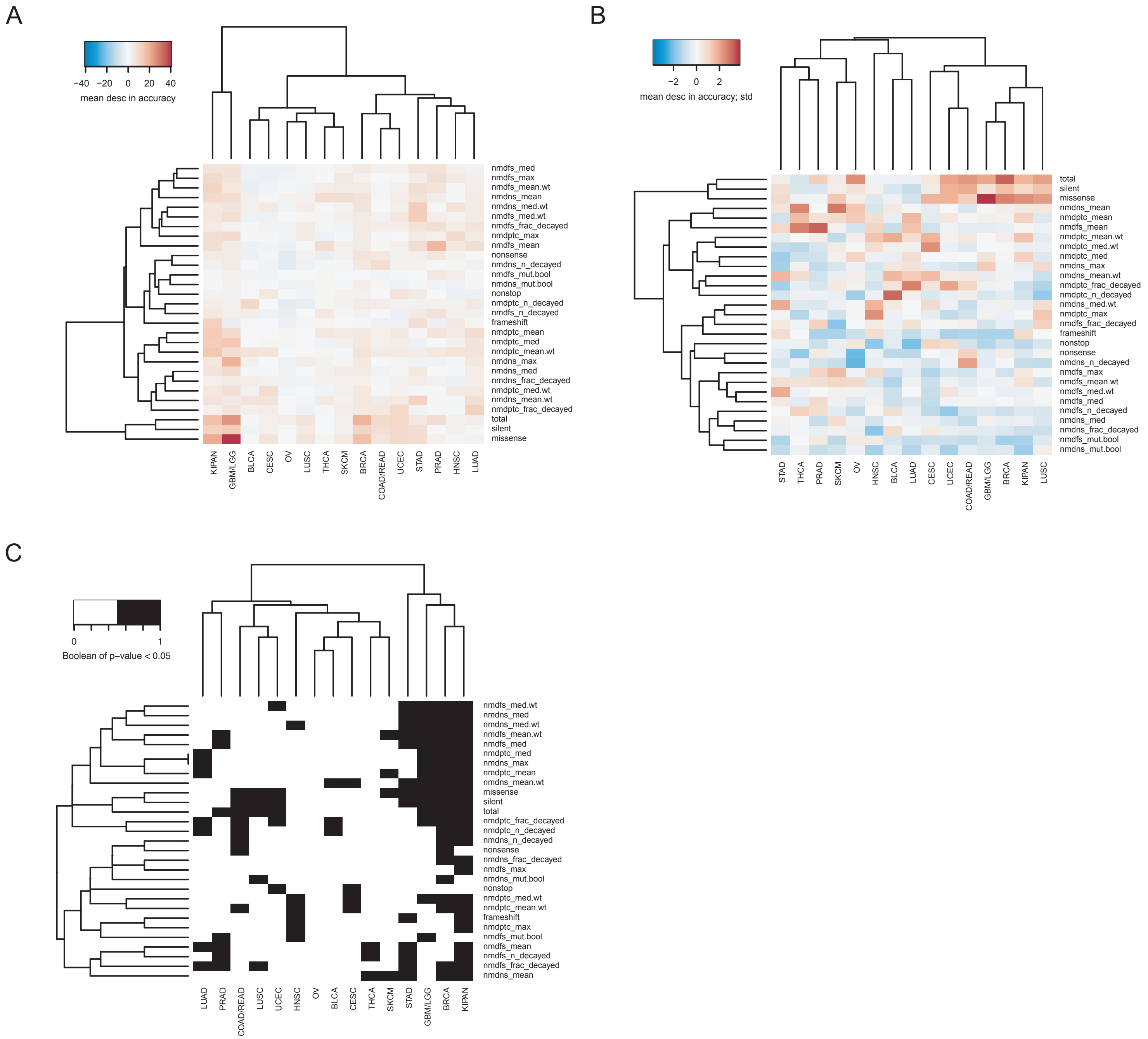

A
−40 −20 0 20 40
mean desc in accuracy
nmdfs_med
nmdfs_max
nmdfs_mean.wt
nmdns_mean
nmdns_med.wt
nmdfs_med.wt
nmdfs_frac_decayed
nmdptc_max
nmdfs_mean
nonsense
nmdns_n_decayed
nmdfs_mut.bool
nmdns_mut.bool
nonstop
nmdptc_n_decayed
nmdfs_n_decayed
frameshift
nmdptc_mean
nmdptc_med
nmdptc_mean.wt
nmdns_max
nmdns_med
nmdns_frac_decayed
nmdptc_med.wt
nmdns_mean.wt
nmdptc_frac_decayed
total
silent
missense
KIPAN
GBM/LGG
BLCA
CESC
OV
LUSC
THCA
SKCM
BRCA
COAD/READ
UCEC
STAD
PRAD
HNSC
LUAD
B
−2 0 2
mean desc in accuracy; std
total
silent
missense
nmdns_mean
nmdptc_mean
nmdfs_mean
nmdptc_mean.wt
nmdptc_med.wt
nmdptc_med
nmdns_max
nmdns_mean.wt
nmdptc_frac_decayed
nmdptc_n_decayed
nmdns_med.wt
nmdptc_max
nmdfs_frac_decayed
frameshift
nonstop
nonsense
nmdns_n_decayed
nmdfs_max
nmdfs_mean.wt
nmdfs_med.wt
nmdfs_med
nmdfs_n_decayed
nmdns_med
nmdns_frac_decayed
nmdfs_mut.bool
nmdns_mut.bool
STAD
THCA
PRAD
SKCM
OV
HNSC
BLCA
LUAD
CESC
UCEC
COAD/READ
GBM/LGG
BRCA
KIPAN
LUSC
C
0 1
Boolean of p−value < 0.05
nmdfs_med.wt
nmdns_med
nmdns_med.wt
nmdfs_mean.wt
nmdfs_med
nmdptc_med
nmdns_max
nmdptc_mean
nmdns_mean.wt
missense
silent
total
nmdptc_frac_decayed
nmdptc_n_decayed
nmdns_n_decayed
nonsense
nmdns_frac_decayed
nmdfs_max
nmdns_mut.bool
nonstop
nmdptc_med.wt
nmdptc_mean.wt
frameshift
nmdptc_max
nmdfs_mut.bool
nmdfs_mean
nmdfs_n_decayed
nmdfs_frac_decayed
nmdns_mean
LUAD
PRAD
COAD/READ
LUSC
UCEC
HNSC
OV
BLCA
CESC
THCA
SKCM
STAD
GBM/LGG
BRCA
KIPAN

Figure S11

A

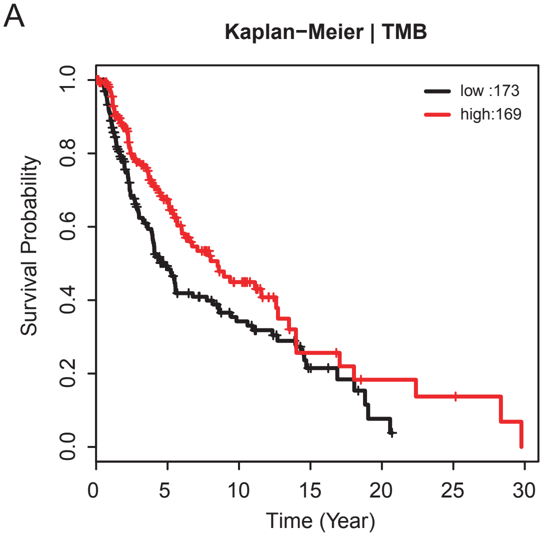


B

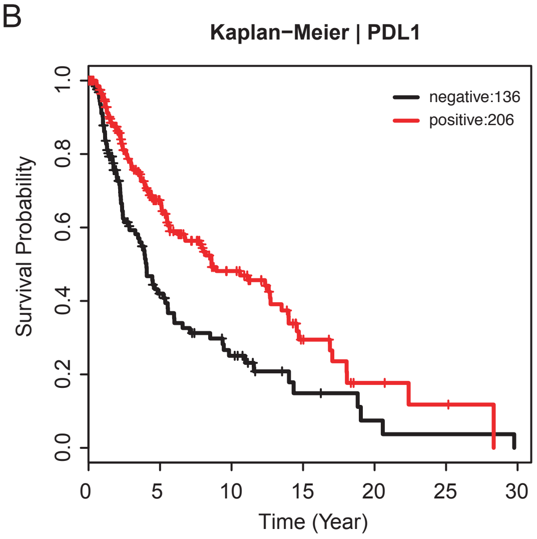
